# Plant secondary metabolite diversity reflects both phylogeny and ecological adaptation

**DOI:** 10.1101/2021.10.26.465835

**Authors:** Simon Pierce, Wen-Yong Guo, Bruno. E.L. Cerabolini, Daniel Negreiros, Franco Faoro, Giulia Magoga, Matteo Montagna, G. Wilson Fernandes, Alberto Spada

**Affiliations:** Department of Agricultural and Environmental Sciences – University of Milan, via Celoria 2, 20133 Milano, Italy; Zhejiang Tiantong Forest Ecosystem National Observation and Research Station, School of Ecological and Environmental Sciences, East China Normal University, 200241 Shanghai, P.R. China; Department of Biotechnologies and Life Sciences (DBSV), University of Insubria, via J.H. Dunant 3, I-21100 Varese, Italy; Ecologia Evolutiva e Biodiversidade/DBG, CP 486, ICB/Universidade Federal de Minas Gerais, 30161-970, Belo Horizonte, MG, Brazil; BAT Center - Interuniversity Center for Studies on Bioinspired Agro-Environmental Technology, University of Napoli “Federico II”, Portici, Italy

**Keywords:** CSR theory, Grime, plant defence/defense, plant functional type, plant toxin, universal adaptive strategy theory

## Abstract

A phylogenetic framework explaining plant secondary metabolite diversity is lacking, but metabolite classes could represent adaptations to habitat resource availability. We test the hypothesis that primary adaptive strategies (competitors, C; stress-tolerators, S; ruderals, R) are associated, respectively, with nitrogenous metabolites synthesized in persistent organs (alkaloids), nitrogen-lacking aromatic terpenes and phenolics, and nitrogenous compounds prevalent in reproductive tissues (cyanogenic glucosides and glucosinolates). A matrix was compiled of 1019 species for which secondary metabolite pathways and CSR strategies are known. Accounting for phylogenetic relatedness and native biomes, we found that most phytochemical pathways did not correlate with strategy axes, but certain key associations were evident. C-selection was positively associated with amino acid-derived phenylpropanoids (low phylogenetic relatedness; λ <0.5) and pyrrolizidine alkaloids and galloyl derivatives (high λ), and negatively with N-lacking linear monoterpenes (low λ). Nitrogenous cyanogenic glucosides positively correlated with R-selection (low λ). Terpenoids were widely distributed, but correlated positively with S- and negatively with R-selection (low λ). Twenty-six correlations between phytochemicals and biomes (low λ) were evident. Most secondary metabolite synthesis pathways are widespread, reflecting common roles and obligate defence, and strong phylogenetic effects are often evident. However, the character of phytochemical/adaptive strategy associations agrees with ecological theory and thus reflects adaptation.

## Introduction

Many plant products and secondary metabolites are perceived as characteristic of specific taxonomic groups, but secondary metabolite diversity is difficult to explain solely in terms of phylogenetic relationships because no overarching phylogenetic pattern is evident^1-7^. Even when specific metabolites are apparently restricted to particular taxa this often reflects a lack of wider investigation and depends on the identity of the species from which the compound was first isolated. For example, Cucurbitacins, typically isolated from the eponymous Cucurbitaceae, are now known to occur throughout a range of disparate families including Elaeocarpaceae, Rosaceae, Rubiaceae, Scrophulariaceae and Sterculiaceae^8-16^. Even the well-studied mustard-oil system (i.e., employing glucosinolates with a myrosinase ‘activator’ compartmentalised in myrosin cells), often viewed as typical of the Brassicaceae, has been recorded from 16 families encompassing nine angiosperm orders and has evolved at least twice in two major ‘mustard-oil’ clades^17^. Cyanogenic glucosides, also often associated with Brassicaceae, are even more widely distributed across major eudicot clades, monocots, gymnosperms and ferns^18,19^. Specific compounds can sometimes be used as chemotaxonomical markers to distinguish taxa, reflecting micro-cladogenic events (e.g., certain iridoid glucosides^20-21^). However, widespread occurrence of secondary metabolite classes is understandable because of their derivation from common precursors in primary metabolism, following minor modification of pre-existing biosynthesis pathways^22^. For example, pathways involved in terpenoid synthesis were apparently present in the ancestor of all land plants^3,5^.

As our view of secondary metabolites shifts from one of firm phylogenetic constraints and taxonomic characters^23^ to one of plants as ‘Jacks-of-all-trades, masters of all’^24^, it is perhaps accurate to say that some taxa tend to favour or emphasise production of certain secondary metabolite classes. Some can specialise in production of very specific molecules within these classes, the result being a ‘patchy’ distribution of secondary metabolites across the plant kingdom^5^. Such apparent trends and specific secondary metabolite roles may reflect a variety of processes, including ancient horizontal gene transfer from bacteria and fungi (and possibly viral recombination), widespread but silent genes, and direct lateral transfer of the metabolites themselves from other organisms, particularly symbiotic endophytes and mycorrhizal fungi^3,5,6^.

The functions and chemical composition of secondary metabolites also potentially help explain their patchy distribution within the plant kingdom. They can be broadly classified as ‘nitrogen containing’ and ‘without nitrogen’^5^, and their production is thus potentially driven by adaptation to different edaphic resource availabilities across habitats, reflecting resource investment trade-offs between growth (involving primary metabolism) and functions such as the attraction of pollinator and seed dispersal vectors and defence against herbivory, pathogens, or abiotic stressors (involving secondary metabolism). Indeed, as primary and secondary metabolism depend largely on the output of the shikimate, polyketide and mevalonic pathways^25^ this potentially leads to competition between primary and secondary metabolism for carbon skeletons and nitrogen. For example, alkaloid synthesis competes with protein synthesis because both use amino acid precursors such as lysine or phenylalanine. The concepts of primary/secondary metabolism trade-offs and of differential allocation to defence at the cost of growth are inherent to various plant defence hypotheses, such as *optimal defence theory*^26^, the *resource availability hypothesis*^27^, and the *growth-differentiation balance hypothesis* (GDBH)^25^.

In experimental studies, the operation of growth/defence trade-offs is suggested by limitation of relative growth rates^28,29^. Trade-offs involving biomass production, flowering, fruiting, and seed production are evident in 82% of controlled studies that investigate defence costs^30^. A recent review^31^ suggests that while growth/defence trade-offs are prevalent across biological systems, they sometimes do not manifest due to allocation to processes additional to growth and defence, such as storage or reproduction. Trade-offs can occur for one class of compounds but not for others^32,33^, and are sometimes only evident under nutrient limiting conditions^34^. Plant cultivation can facilitate both growth and defence, masking the fitness value of trade-offs operating in the wild^31^. Thus, allocation trade-offs are evidently complex and depend on the simultaneous action of multiple factors, both internal and external^1^, which has led to difficulty placing the cost of secondary metabolite production in the context of overall plant fitness, although the absolute energetic cost of metabolite production is likely to be significant^5^.

In light of the complexity and context-dependency of growth/defence trade-offs and secondary metabolite production, is a general explanation of trends in secondary metabolite production possible?

A general explanation of plant functioning is attempted by plant ecological strategy theories. A “strategy” is the regime of resource investments between numerous adaptive traits in the face of multiple natural selection pressures^36^, or “an instantaneous process of resource allocation between competing functions, maximising fitness across contrasting niches during development”^37^. Secondary metabolites are potentially functional traits (i.e., characters that affect fitness and survival^38^), with interspecific variability in defence metabolites and defence/herbivory relationships playing a significant role in species coexistence^39^. The concept of ‘plant defence syndromes’ comprising a range of traits and multiple defence compounds^4,24^ is consistent with that of plant ecological strategies, and indeed can be conceptualized as a component of the general plant strategy. The principal components of plant functional variability world-wide are summarized by the ‘global spectrum of plant form and function’: an observation of the importance of plant resource economics (acquisitive *vs.* conservative management of matter and energy) and stature (organ and whole-plant size)^40^. The resource economics spectrum essentially represents a nitrogen allocation trade-off between structural (cell wall) investment and metabolism^41^, highlighting the competing roles of mineral resources in plant structural and metabolic adaptations and the potential integration of trade-offs in both primary and secondary functioning, and does appear to be involved in plant defence trade-offs^42,43^.

Currently the only theory that can explain resource economics and stature as the principal evolutionary responses to environmental selection pressures is Grime’s^44-46^ competitor, stress-tolerator, ruderal (CSR) theory, for which economics and size traits delimit a common frame of reference across vascular plant life forms^47-51^. Notably, CSR theory was cited by, and thus evidently inspired, subsequent growth/defence trade-off theories^25,27^, and thus CSR theory promises to provide a general plant eco-evolutionary context in which to understand adaptive/defence specialisations.

Which specific specialisations can we expect?

S-selected ‘stress-tolerators’, no matter what type of stressor or limitation to metabolic performance, tend to invest in constitutive physical defence (inherently tough and persistent tissues) forming part of a conservative strategy in which resources and biomass are gradually accrued^52,53^. S-selection is thus associated with high tissue dry matter content, high C:N, and a greater extent of mechanical and parenchyma tissues^47,54^ which probably reflects greater investment in cell wall materials and reserves that can be used to support metabolism during periods of limitation. The secondary metabolites associated with S-selection could thus be expected to respect this high C:N and be based predominantly on compounds lacking N, such as phenolics and terpenoids.

Alternatively, C- and R-selected species occupy relatively nutrient-rich habitats and can be expected to experience greater availabilities of mineral resources and thus N-containing secondary metabolites. Alkaloids are typically synthesized in roots or rhizomes and are subsequently translocated to and accumulated in other plant parts^55-57^. Occasionally alkaloid biosynthesis occurs in leaves (e.g., caffeine accumulates in seeds of *Coffea arabica* from biosynthesis in younger leaves^58^) or in stem sieve elements adjacent to laticifers^59^, but rarely does alkaloid synthesis occur in fruits and seeds. Thus, alkaloid production is more prevalent in larger perennial herbs with persistent root systems and rhizomes, and can be expected to be associated with C-selection. N-containing compounds such as cyanogenic glucosides and glucosinolates can be synthesized in any plant part but are typically found in elevated concentrations in reproductive structures, particularly fruits and seeds, for both annual^60,61^ and perennial^62^ species. This emphasis on the support of reproductive function is also a key concept for R-selection, and we can reasonably expect a tendency towards cyanogenic glucoside and glucosinolate production in relatively R-selected species. However, exceptions to these possible trends exist (e.g., alkaloids in the annual opium poppy, *Papaver somniferum*, cyanogenic glucosides in rosaceous stone fruits), and the extent to which these different types of secondary metabolite defences are embroiled in primary adaptive strategies remains to be tested. This was the main objective of the present study.

Specifically, our hypotheses were that: 1). the extent of S-selection is correlated with production of secondary metabolites based on N-lacking molecules, such as terpenoids or phenolics, 2). the degree of C-selection is associated with capacity to synthesize N-containing alkaloids, 3). R-selection is associated with N-containing secondary metabolites typically found in greater quantities in reproductive tissues, such as cyanogenic glucosides and glucosinolates. Additionally, we expect these compounds to reflect the environment and thus be differentially represented in distinct biomes. Here, we compare the extent of C-, S- and R-selection with the ability to synthesize different classes of phytochemicals evident from a global database of vascular plant species for which both CSR strategies and secondary metabolite compounds are known. We use phylogenetic linear regression to account for phylogenetic relatedness, and for relationships between phytochemical clusters and biomes world-wide. While this approach may include limitations (such as not being able to differentiate between obligate or facultative use of secondary metabolites, and restriction to a small portion of the world flora) it is presently the most comprehensive way of placing secondary metabolites in a general ecological context, using the largest globally relevant dataset currently available.

## Materials and methods

CSR strategies for thousands of plant species world-wide are available as supplementary material to the global CSR strategy analysis of Pierce *et al*.^63^, which was employed here. For each species, we searched the global phytochemical interactions database (PCIDB; www.genome.jp/db/pcidb) and found a total of 1019 vascular plant species present in both databases, encompassing 52 orders and 139 families of vascular plants, and including 892 dicotyledonous species (basal and eudicots), 87 monocotyledons, 19 gymnosperms and 21 pteridophytes. Thus, while each separate database includes many thousands of species, the current study was restricted to the species present in both databases (i.e., not all species known to produce, for example, glucosinolates, were included and not all species known to be, for instance, strongly R-selected were present).

The PCIDB has the particular advantage of a standardised classification system of secondary metabolites, employing 83 phytochemical cluster classes which include information on chemical structure, including KNApSAcK metabolite codes^64^ and KEGG Chemical Function-and-Substructures codes (KCF-S)^65^ (the full list of clusters is presented in Table S1[**Supplementary Information**]). Species in the PCIDB are classified according to their National Center for Biotechnology Information (NCBI) taxonomic name and ID code (www.ncbi.nlm.nih.gov/taxonomy)^66^, and in the present study the NCBI open chemistry database (https://pubchem.ncbi.nlm.nih.gov) was also referred to in order to double-check chemical formulas and structures, specifically with regard to N-containing *vs.* N-lacking compounds. Species nomenclature was checked according to the Plant List (www.theplantlist.org) to account for synonyms between NCBI and the nomenclature used by the CSR strategy database of Pierce *et al*.^63^.

Associations between secondary metabolite use and the native biome(s) of each species were also investigated. Fourteen biomes were defined according to Olson *et al*.^67^. Information regarding the biome(s) within the native range of each species was obtained from the comparison of the global distribution of biomes with the georeferenced records of each species (obtained from GBIF; www.gbif.org/species/search) occurring inside its native range. The native range of each species was obtained from sources such as GRIN taxonomy for plants (www.ars-grin.gov/cgi-bin/npgs/html/tax_search.pl?language=en), eMonocot (http://emonocot.org) and Catalogue of Life (www.catalogueoflife.org/col/search/all). The C-, S-, and R-values of species, the number of compounds in each phytochemical cluster known for each species, and data regarding presence/absence in biomes were collated into a matrix.

As leaf economics traits are subject to phylogenetic effects^68^, phylogenetic relatedness was accounted for by constructing a phylogenetic tree for the study species based on the dated supertree of Zanne *et al.*^69^, corrected and extended by a recent tree including over 31,000 species based on several genetic markers^70^. All analyses were performed in the R environment^71^. An R function, *S.PhyloMaker*^71^ was used to produce phylogenies for subsets of species using Qian and Jin’s^70^ “Scenario 3” approach (i.e., adding absent species to their families or genera following Phylomatic and BLADJ^72^). Phylogenetic linear regression was applied to analyse the relationships between the 83 clusters, C-, S- and R-values and biome classes while accounting for phylogenetic relatedness, using the R package *phylolm*^73^. The phylogenetically corrected coefficient of correlation was denoted by *α* (using the *p* ≤ 0.05 level), and the strength of phylogenetic relatedness was denoted by λ (on a scale of 0 to 1, with λ <0.5 interpreted as relatively low phylogenetic relatedness).

## Results

After phylogenetic correction, only seven out of 83 clusters were significantly correlated with C-, S- or R-selection, and only five of these with relatively low phylogenetic relatedness between species (λ <0.5). The extent of C-selection was significantly positively correlated with the production of pyrrolizidine alkaloids (cluster 2: *α* = 0.099, *p* = 0.004) and galloyl derivatives and ellagitannins (cluster 81: *α* = 0.062, *p* = 0.015) but with strong phylogenetic relatedness (λ = 0.782 and 0.853, respectively) (Fig. 1A,D). C-selection was significantly positively correlated with coumarin and furanocoumarin phenylpropanoids (cluster 25; *α* = 0.092, *p* = 0.036) with a low phylogenetic effect (λ = 0.395; Fig. 1B), and negatively correlated with linear monoterpenes (cluster 34: *α* = -0.079, *p* = 0.004), with a negligible phylogenetic effect (λ = 0.000; Fig. 1C). The ten species in the database that produced the greatest number of pyrrolizidine alkaloids were all classed as C, C/CR, C/CSR or CR-selected: for example, *Symphytum officinale* (tertiary CSR strategy = C/CR; C:S:R = 69.7 : 0.0 : 30.3 %) is known to produce eight distinct pyrrolizidine alkaloids. The species exhibiting the greatest diversity in aromatic coumarin and furanocoumarin phenylpropanoids were the C-selected Apiaceae *Heracleum sphondylium* (C:S:R = 82.8 : 6.8 : 10.5 %), *Angelica archangelica* (C:S:R = 81.4 : 8.8 : 9.8 %) and *Peucedanum ostruthium* (C:S:R = 66.2 : 17.1 : 16.7 %).

**Figure 1.**
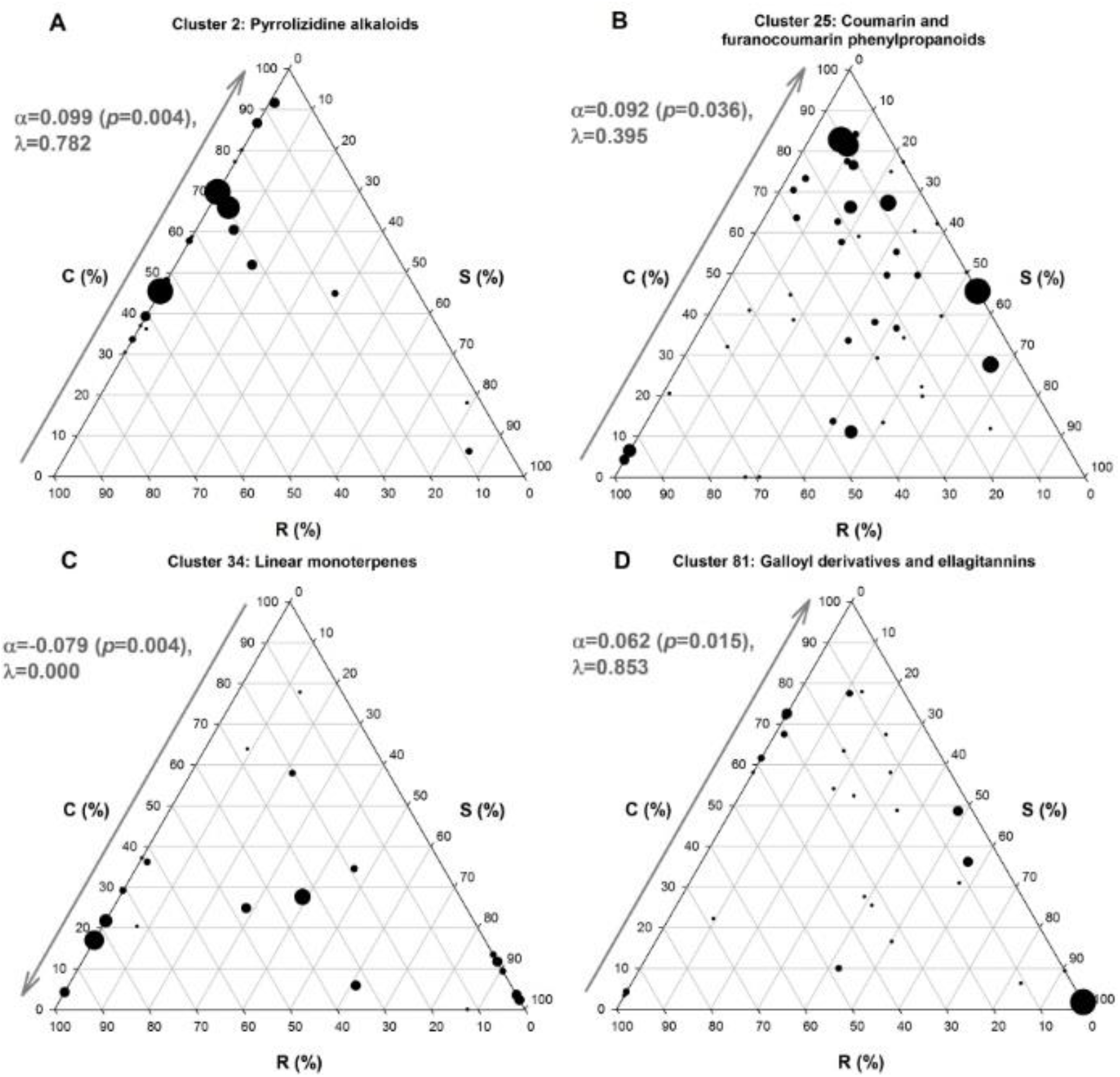
Significant phylogenetically corrected correlations (*α*; positive and negative) between the extent of C-selection (i.e., a predominantly competitor adaptive strategy) and the number of species producing secondary metabolites belonging to different phytochemical clusters, with the strength of phylogenetic relatedness between species (λ) indicated (1.000=highest relatedness, 0.000=lowest relatedness). Clusters (as detailed in Table S1) are: A). pyrrolizidine alkaloids (cluster 2), B). coumarin and furanocoumarin phenylpropanoids (cluster 25), C). linear monoterpenes (cluster 34), and D). galloyl derivatives and ellagitannins (cluster 81). The size of points is proportional to the number of compounds of each class produced per species.

The extent of S-selection was positively correlated with production of terpenoids (cluster 38; *α* = 0.287, *p* = 0.045), with low phylogenetic relatedness (λ = 0.399; Fig. 2A), and with cadinanes (cluster 39; *α* = 0.044, *p* = 0.041, λ = 0.759; Fig. 2B) and labdanes (cluster 46; *α* = 0.019, *p* = 0.038, λ = 0.235; Fig. 2C). Species that produced ten or more different kinds of terpenoids included the strongly S-selected *Cistus creticus* (C:S:R = 11.8 : 88.2 : 0.0 %)*, Acorus calamus* (C:S:R = 5.8 : 61.1 : 33.1 %)*, Rosmarinus officinalis* (C:S:R = 3.5 : 96.5 : 0.0 %) and *Pinus halepensis* (C:S:R = 2.4 : 97.6 : 0.0 %), but also some C-selected species (e.g., *Valeriana officinalis*; C:S:R = 80.4 : 0.0 : 19.6 %) and broadly R-selected species (e.g., *Artemisia annua*; C:S:R = 24.9 : 28.3 : 46.8 %). The S-selected species mentioned here also produced the greatest variety of cadinanes. Species producing the greatest numbers of labdanes included, again, S-selected angiosperms such as *Cistus creticus* and *Marrubium vulgare* (C:S:R = 15.9 : 66.8 : 17.3 %), extremely S-selected gymnosperms (e.g., *Platycladus orientalis*, C:S:R = 0.0 : 100.0 : 0.0 %; *Juniperus communis*, C:S:R = 0.0 : 86.6 : 13.4 %), but also some domesticated cereals such as *Zea mays* (C:S:R = 83.0 : 8.6 : 8.4 %) and *Oryza sativa* (C:S:R = 37.9 : 37.1 : 25.0 %).

**Figure 2.**
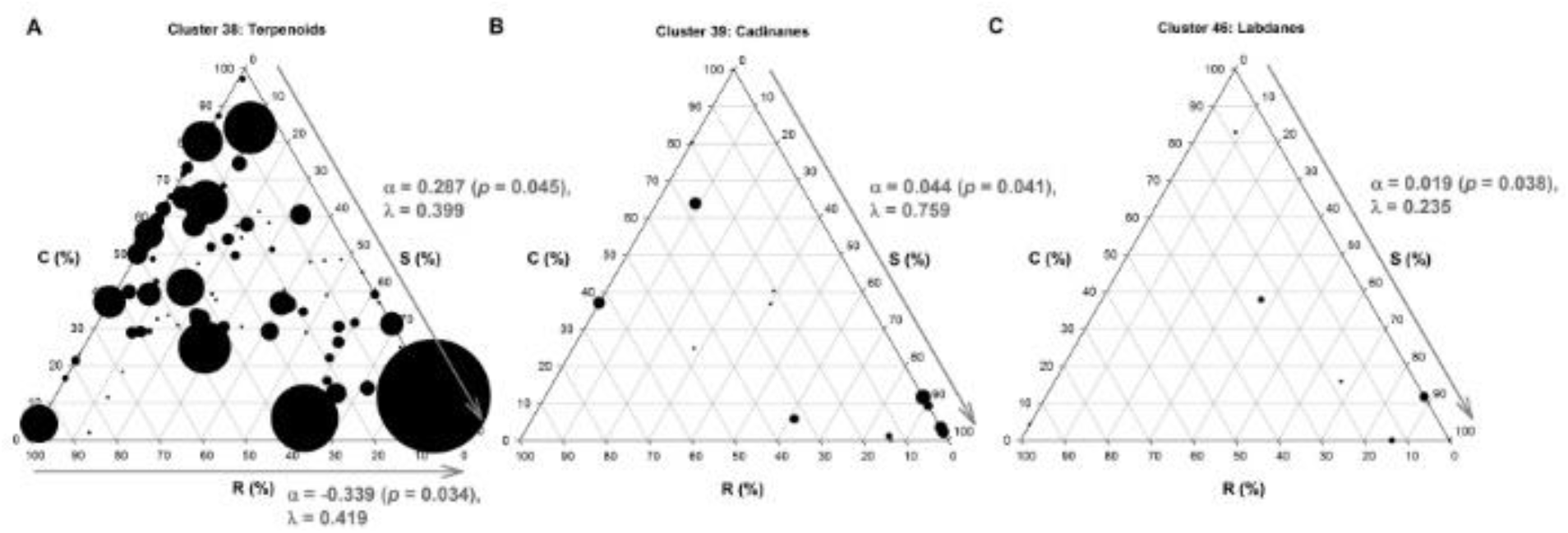
Significant phylogenetically corrected correlations (*α*; positive and negative) between the extent of S-selection and the number of species producing secondary metabolites belonging to different phytochemical clusters, with the strength of phylogenetic relatedness between species (λ) indicated (1.000=highest relatedness, 0.000=lowest relatedness). Clusters (as detailed in Table S1) are: A). terpenoids (cluster 38), B). cadinanes (cluster 39), and C). labdanes (cluster 46). The size of points is proportional to the number of compounds of each class produced per species.

The extent of R-selection was negatively correlated with flavanone flavonoids (cluster 14; *α* = -0.127, *p* = 0.047, λ = 0.448; Fig. 3A) and terpenoids (cluster 38; *α* = -0.339, *p* = 0.034, λ = 0.419; Fig. 2A), but positively correlated with production of cyanogenic glucosides derived from valine or isoleucine (cluster 73; *α* = 0.032, *p* = 0.029, λ = 0.000; Fig. 3B). Species exhibiting production of multiple flavanone flavonoids included a wide range of perennial species encompassing C-selected woody angiosperms (*Paulownia tomentosa,* C:S:R = 84.2 : 2.7 : 13.2 %; *Vitis vinifera,* C:S:R = 72.5 : 0.0 : 27.5 %), S-selected gymnosperms (*Cedrus deodara,* C:S:R = 0.9 : 99.1 : 0.0 %), a range of both woody and herbaceous Fabaceae exhibiting intermediate CSR strategies (*Amorpha fruticosa,* C:S:R = 40.4 : 39.1 : 20.5 %; *Lupinus luteus,* C:S:R = 58.5 : 0.0 : 41.5 %; *Phaseolus vulgaris,* C:S:R = 57.7 : 19.4 : 22.9 %; *Robinia pseudoacacia,* C:S:R = 47.9 : 41.4 : 10.7 %), but also some R-selected species such as *Arabidopsis thaliana* (C:S:R = 4.3 : 0.0 : 95.7 %). The ten species exhibiting the greatest diversity of cyanogenic glucosides included extremely R-selected *Linum usitatissimum* (Linaceae; Fabid eudicot clade; C:S:R = 1.9 : 12.9 : 85.2 %) and *Arabidopsis thaliana* (Brassicaceae; Malvid eudicot clade), R-selected herbaceous Fabaceae such as *Trifolium repens* (C:S:R = 36.2 : 1.6 : 62.2 %)*, Lotus corniculatus* (C:S:R = 14.2 : 19.0 : 66.8 %)*, Ornithopus perpusillus* (C:S:R = 8.0 : 23.7 : 68.4 %)*, Lotus maritimus* (C:S:R = 23.8 : 30.8 : 45.4 %), and CR-selected Ranunculaceae (i.e., basal eudicots) such as the forb *Caltha palustris* (C:S:R = 59.1 : 0.0 : 40.9 %) and the vine *Clematis flammula* (C:S:R = 41.1 : 24.5 : 34.4 %). However, tree species were also able to produce cyanogenic glucosides, such as *Euonymus japonicus* (S/CS), *Fagus sylvatica* (S/CS) and *Robinia pseudacacia* (CS).

**Figure 3.**
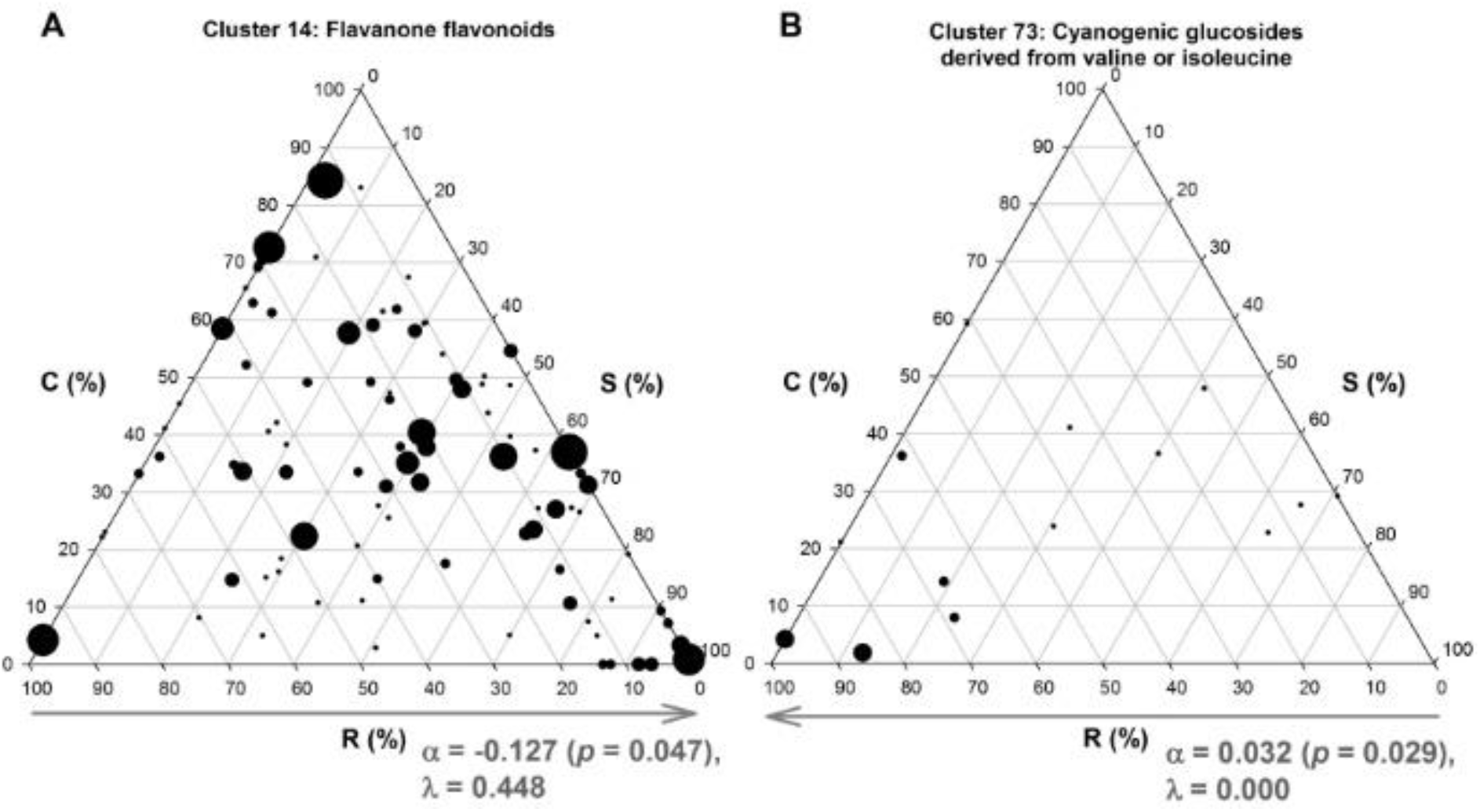
Significant phylogenetically corrected correlations (*α*; positive and negative) between the extent of R-selection and the number of species producing secondary metabolites belonging to different phytochemical clusters, with the strength of phylogenetic relatedness between species (λ) indicated (1.000=highest relatedness, 0.000=lowest relatedness). Clusters (as detailed in Table S1) are: A). flavanone flavonoids (cluster 14), and B). cyanogenic glucosides derived from valine or isoleucine (cluster 73). The size of points is proportional to the number of compounds of each class produced per species.

Twenty-six significant correlations between clusters and global biomes were evident with low phylogenetic relatedness (Table 1). Notably, N-lacking compounds such as coumestanes and neoflavonoids (cluster 17) and aphidicolane terpenoids (cluster 41) were positively correlated with deserts and xeric shrublands (*α* = 0.145, *p* = 0.003 and *α* = 0.530, *p* = 0.012, respectively; Table 1A). The biome associated with the greatest phytochemical diversity, with low phylogenetic effects, was that of montane grasslands and shrublands. These were positively associated with diversity in caffeate and tyramine derivative alkaloids (cluster 6), steroid alkaloids (cluster 11), lignans (cluster 21), terpenoids (cluster 38), carotenoids (cluster 59) and galloyl derivatives/ellagitannins (cluster 81) (see Table 1A for respective *α* and *p* values). A range of clusters were significantly negatively correlated with temperate forests: purine alkaloids (cluster 12) with temperate broadleaf/mixed forest, while cadinanes (cluster 39), aphidicolane terpenoids (cluster 41) and cyanogenic glucosides derived from valine or isoleucine (cluster 73) correlated negatively with temperate coniferous forests (Table 1A). Tropical and subtropical moist broadleaf forests were particularly associated with diversity in iridoids (cluster 36), and negatively with pyrrolidine alkaloids (cluster 1) (Table 1A). Boreal forests and taiga, Mediterranean forests, woodlands and scrub, and tundra biomes were not significantly correlated with any particular phytochemical cluster (Table 1B).

**Table 1.** Phylogenetic linear regression: significant correlations (*α*; both positive and negative) between biomes (as defined by Olson *et al.,* 2001) and phytochemical clusters after phylogenetic correction (A) with an indication of biomes for which no correlations were evident (B).

| **(A). Biomes exhibiting significant correlations** | **Phytochemical clusters** | **$\alpha$** | **$p$** |
| --- | --- | --- | --- |
| Deserts and Xeric Shrublands | 17 Coumestanes, | 0.1448 | 0.003 |
|  | 41 Terpenoids | 0.5297 | 0.012 |
| Flooded Grasslands Savannas | 36 Iridoids | 0.3168 | 0.046 |
| Mangroves | 13 Chalcones, | 0.5977 | 0.004 |
|  | 30 Stilbenoid dimers, stilbenoid oligomers and bis(bibenzyl)s | 0.3527 | <0.001 |
| Montane Grasslands Shrublands | 6 Caffeate derivative alkaloids, | 0.2789 | 0.015 |
|  | 11 Steroid alkaloids, | 0.6325 | 0.001 |
|  | 21 Lignans, | 0.1699 | 0.002 |
|  | 38 Terpenoids, | 0.4541 | 0.027 |
|  | 59 Carotenoids, | 0.1925 | 0.004 |
|  | 81 Galloyl derivatives and ellagitannins | 0.0808 | 0.006 |
| Temperate Broadleaf and Mixed Forests | 12 Purine alkaloids, | -0.0520 | 0.009 |
|  | 59 Carotenoids | -0.1667 | 0.041 |
| Temperate Coniferous Forests | 39 Cadinane terpenoids, | -0.0701 | 0.011 |
|  | 41 Aphidicolane terpenoids, | -0.7319 | <0.001 |
|  | 73 Cyanogenic glucosides derived from valine or isoleucine | -0.0486 | 0.016 |
| Temperate Grasslands Savannas and Shrublands | 14 Flavanone flavonoids, | 0.1748 | 0.033 |
|  | 77 Glucosinolates group B | -0.1226 | 0.008 |
| Tropical and Subtropical Coniferous Forests | 17 Coumestanes | -0.2498 | <0.001 |
| Tropical and Subtropical Dry Broadleaf Forests | 4 Isoquinoline alkaloids, | -0.3231 | 0.011 |
|  | 17 Coumestanes, | 0.2311 | 0.006 |
|  | 71 Betacyanins and betaxanthins | 0.0259 | 0.016 |
| Tropical and Subtropical Grasslands Savannas and Shrublands | 21 Lignans | 0.1658 | 0.037 |
| Tropical and Subtropical Moist Broadleaf Forests | 1 Pyrrolidine alkaloids, | -0.1575 | 0.043 |
|  | 36 Iridoids, | 0.1862 | 0.032 |
|  | 71 Betacyanins and betaxanthins | 0.0166 | 0.024 |
| **(B). Biomes not correlated with any phytochemical cluster** |  |  |  |
| Boreal Forests Taiga | None |  | >0.05 |
| Mediterranean Forests Woodlands Scrub | None |  | >0.05 |
| Tundra | None |  | >0.05 |

## Discussion

Analysis of a global dataset demonstrates that while most secondary metabolites are widespread (reflecting roles as essential pigments, membrane components and general constitutive defences) and strong phylogenetic effects are often evident, the distribution of some secondary metabolites is associated with primary ecological strategies and native biomes. In these cases, the specific trends are also consistent with explanations suggested by ecological theory^44-46,53^. In extreme summary: 1). certain nitrogenous compounds synthesized mainly in perennial plant parts (or some aromatics also derived from amino acid synthesis) are correlated with C-selection and thus adaptation to resource-rich, stable habitats, 2). some N-lacking aromatic compounds are associated with S-selection and thus adaptation to resource-poor habitats, and 3). N-containing cyanogenic glucosides are associated with R-selection and thus adaptation to resource-rich but unstable habitats. This statement is extremely general and the correlations, while statistically significant, are imperfect (e.g., not all competitors produce pyrrolizidine alkaloids, while some stress-tolerators do; Fig. 1A), but this can potentially offer a general explanation as to why certain groups of secondary metabolites seem to be favoured by particular species: species that share a general approach to survival can share a general approach to defence, even when they are not closely related phylogenetically.

Specifically, hypothesis 1 suggested that the extent of S-selection is correlated with production of secondary metabolites based on N-lacking molecules, such as terpenoids or phenolics. Positive correlations with the production of terpenoids, cadinanes and labdanes bears this out. The character of the species producing these secondary metabolites also agrees with the hypothesis: most are strongly S-selected perennial species and represent a range of gymnosperms and angiosperms, including a number of Mediterranean taxa (e.g., *Cistus creticus, Marrubium vulgare, Pinus halepensis, Rosmarinus officinalis*). The positive association of the desert and xeric shrubland biome with N-lacking coumestanes and aphidicolanes (but not with alkaloids or other N-containing secondary metabolites) also agrees with this hypothesis.

Hypothesis 2 suggested that the degree of C-selection is associated with capacity to synthesize N-containing alkaloids, which are usually produced in perennial organs. The findings were broadly supportive of the hypothesis but outlined a slightly more complicated picture. The positive correlation of C-selection with pyrrolizidine alkaloids suggests that these may reflect adaptation to productive habitats, but some classes of alkaloids appear to be extremely widespread (pyrrolidine alkaloids, isoquinoline alkaloids). C-selection was also associated with production of certain aromatic compounds, particularly coumarin and furanocoumarin phenylpropanoids and galloyl derivatives. Many C-selected Apiaceae, such as *Cicuta virosa, Chaerophyllum hirsutum* and *Oenanthe fistulosa*, also produced polyynes, as did numerous CS- to CR/CSR-selected Fabaceae and Araliaceae. C-selected species producing coumarin and furanocoumarin phenylpropanoids were large Apiaceae such as *Angelica archangelica, Heracleum sphondylium* and *Pastinaca sativa*, Asteraceae such as *Inula helenium*, and other broadly C/CS- or C/CR-selected members of Solanaceae (*Atropa belladonna*), Calophyllaceae (*Calophyllum brasiliense*), Sapindaceae (*Aesculus hippocastanum*) and Moraceae (*Ficus* spp., *Morus* spp.). Crucially, although the final coumarin and furanocoumarin phenylpropanoid products are N-lacking aromatic compounds, they are derived from amino acid synthesis pathways that rely on N availability. Polyynes are also relatively complex molecules characterised by carbon chains with alternating triple bonds, and often including multiple aromatic sub-groups. Thus, it appears that C-selection is not necessarily associated with N incorporated directly in the final defence compound *per se*, but with N availability during synthesis.

The third hypothesis, that R-selection is associated with nitrogenous secondary metabolites typically used to protect reproductive structures, such as cyanogenic glucosides and glucosinolates, was found to be supported in the case of cyanogenic glucosides only. Cyanogenic glucosides are associated with R-selection across a range of families including Brassicaceae, Fabaceae, Linaceae, Phytolaccaceae, Ranunculaceae, Resedaceae and Tropeolaceae, with negligible phylogenetic effects (Fig. 3). Some CS- and S/SC-selected trees of various families, notably Fabaceae, Rosaceae and Myrtaceae were also able to produce cyanogenic glucosides, in agreement with the known production of these compounds in genera such as *Acacia, Prunus* and *Eucalyptus*, respectively^19,62^.

These associations and correlations may not be perfect, but imperfect correlations are an expected outcome of natural variability. Additionally, some of this variability may not be entirely natural. Many of the notable exceptions, such as relatively C-selected cereals that produce a range of aromatic labdanes (*Zea mays* and *Oryza sativa*) or the excessively chalcone-rich *Vitis vinifera*, are domesticated species. These could embody Züst and Agrawal’s^31^ observation that cultivation alters the pressures selecting for specific growth/defence trade-offs with respect to the wild. Indeed, in the present study no wild graminoids produced labdanes, and Schmelz *et al.*^74^ confirm that these are known within Poaceae only from domesticated cereals. *Vitis vinifera* has been artificially selected for a rich palate of organoleptic qualities that reflect, in part, secondary metabolite diversity. Indeed, domestication of *V. vinifera* involved selection of a range of genes affecting berry development, sugar transport and secondary metabolite production, especially monoterpenoid indole alkaloid biosynthesis, proanthocyanidin accumulation and stilbenoid biosynthesis^75^. Domestication is a process of adaptation to symbiosis with the human species, in which some of the work (notably plant defence) is carried out by the human symbiont and for which the plant is exposed to fewer natural selection pressures^6,76,77^. Thus, traits such as secondary metabolites that are strongly selected for in the wild can become less decisive for survival under cultivation and may be prone to genetic drift or artificial selection. From a wild situation in which plants maintain lasting resistance using a complex mixture of defence compounds, domestication involves human preference for monoculture and small numbers of externally applied defence compounds, potentially leading to rapid herbivore and pathogen adaptation^6^. Experimental studies have found that, across a range of cultivated taxa, domestication does alter plant defences and rates of herbivory, although changes in growth/defence allocation trade-offs are inconsistent and depend on the plant/herbivore system^78^. There is unlikely to be a simple ‘domestication = less innate defence’ relationship as each species has a different propensity to allocate resources to additional functions such as storage or stress resistance^78^. Nonetheless, the original ecological strategies of wild ancestral species are likely to influence defensive capacity following domestication.

Although some specific correlations between phytochemical clusters and biomes were found (such as the relationship between certain N-lacking metabolites and the desert biome) no overarching general trend emerged, probably reflecting the fact that each biome can include a wide range of habitats differing in productivity, causing ‘noise’ in the analysis. Indeed, biome/CSR strategy associations have been observed, but are highly variable for this reason^63^. It is likely that comparison of specific habitats and the presence of species synthesizing distinct phytochemicals would yield more specific insights regarding the relationship between strategies, secondary metabolites and environmental resource availabilities. In the present study, complications also arose from the highly generalised nature of the available phytochemical data (i.e., simply the presence/absence of compounds recorded for each taxon, revealing little of whether metabolites are constitutive or induced), lack of information regarding the site of synthesis and/or accumulation, and the relative energetic costs of each defence mechanism. Indeed, secondary metabolite contents may increase in response to abiotic stresses^79^, and the current dataset can reveal nothing of responses or absolute concentrations of metabolites. Again, it could be hypothesized that metabolite concentration and the relative energetic cost of each metabolite pathway is related to habitat and adaptive strategy. Unfortunately, due to the complexity of metabolite synthesis pathways, currently a precise indication of the synthesis costs of major secondary metabolite classes is unachievable^35^. Data in the present study are also extremely general in the sense that they represent mixed records – i.e., detections of distinct compounds in different samples by different laboratories – for ‘the species’. Almost nothing is currently known of the variability of the secondary metabolite garniture and concentrations of each compound between individuals within populations, and the extent to which this is genetically determined or reflects responses to soil nutrient availabilities.

As with other plant functional traits, intra-specific diversity in secondary metabolites and population-level variability is a prerequisite for natural selection, and almost certainly occurs. Thus, at fine scales, such as within habitats or along environmental gradients of productivity or stress, it is likely that growth/defence trade-offs occur, are important for the structuring of plant communities, and involve strategy-dependent differences in phytochemical production between species and between individuals within populations. Indeed, just as plant species diversity is tightly correlated with strategy diversity and functional trait variance^80^, phytochemical diversity is likely to have been extended via the same eco-evolutionary feedbacks (*sensu* Grime and Pierce^53^). Hopefully, using the applied CSR analysis method used here, we can construct an evolutionary and ecological context for plant secondary metabolites as functional traits that contribute to fitness when plants live together and with other organisms.

In conclusion, most secondary metabolite synthesis pathways are widely evident, reflecting common roles as membrane components, pigments, and general obligate defences. However, additional secondary metabolic specializations appear to have emerged as an integral part of plant primary adaptive strategies in response to fundamental constraints of habitat productivity and stability. This represents further evidence in agreement with the concepts of secondary metabolites as functional characters affecting fitness, and of secondary metabolite involvement in complex growth/defence trade-offs occurring during plant adaptation.

## Acknowledgements

This project was funded with the aid of grant number 19628 (RV_RIC_AT16SPIER; *Piano di Sostegno alla Ricerca* 2015-2017) awarded to S.P. from the Department of Agricultural and Environmental Sciences (DiSAA), University of Milan, Italy.

## Author contributions

S.P., A.S., F.F. and B.E.L.C. conceived the work and compiled the database. S.P. also coordinated between work groups and wrote the manuscript. G.M., M.M. and W.-Y.G. performed the phylogenetic logistic regressions, D.N. and G.W.F. analysed associations between secondary metabolite pathways and biomes.

