## Supplementary material for "Plant secondary metabolite diversity reflects both phylogeny and ecological adaptation": Table S1

**Table S1.** Phytochemical clusters of the global phytochemical interactions database (PCIDB; www.genome.jp/db/pcidb/kna_clst7s) referred to in the present study, including the number of KEGG Chemical Function-and-Substructures (KCF-S) clusters (Kotera *et al.*, 2013) and KEGG BRITE phytochemical compound classes (www.genome.jp/kegg/brite.html) within each general phytochemical cluster.

| Phytochemical cluster | Number of KCF-S clusters | Classes of Phytochemical Compounds in KEGG BRITE |
|---|---|---|
| 1 | 59 | Pyrrolidine alkaloids<br>Tropane alkaloids<br>Piperidine alkaloids<br>Pyridine alkaloids<br>Acetate derived alkaloids |
| 2 | 20 | Pyrrolizidine alkaloids<br>Indolizidine alkaloids |
| 3 | 16 | Quinolizidine alkaloids |
| 4 | 95 | Isoquinoline alkaloids<br>Indole alkaloids<br>Pyrroloindole alkaloids<br>Others |
| 5 | 2 | Thiazole alkaloids |
| 6 | 22 | Tyramine derivatives<br>Caffeate derivatives<br>Coniferyl alcohol derivatives<br>Paracoumaryl alcohol derivatives<br>Sinapate derivatives<br>Others |
| 7 | 42 | Acridone alkaloids<br>Quinazoline alkaloids<br>Quinoline alkaloids |
| 8 | 2 | Imidazole alkaloids |
| 9 | 2 | Phenylalanine derived alkaloids |
| 10 | 20 | Terpenoid alkaloids |
| 11 | 27 | Steroid alkaloids<br>Cholestane<br>Spirostan<br>Furostan<br>Ergostane<br>Stigmastane<br>Cycloartane |
| 12 | 4 | Purine alkaloids |
| 13 | 18 | Aurones<br>Chalcones<br>Dihydrochalcones<br>Stilbenes |
| 14 | 20 | Flavanones<br>Dihyroflavonols<br>Flavan 3-ols<br>Flavan 4-ols<br>Flavan 3,4-diols<br>Isoflavanones |
| 15 | 54 | Flavones<br>Flavonols<br>Flavans<br>Anthocyanidins and anthocyanins<br>Isoflavanes<br>Isoflavones<br>Pterocarpanes<br>Rotenones<br>2-Arylbenzofurans<br>Chromones<br>Xanthenes |
| 16 | 1 | 3-Arylcoumarins |
| 17 | 5 | Coumestanes<br>Neoflavonoids |
| 18 | 3 | Biflavonoids and polyflavonoids |
| 19 | 4 | Proanthocyanidins |
| 20 | 1 | Diels-Alder adduct of chalcone |

| | | |
|---|---|---|
| 21 | 19 | Lignans |
| 22 | 4 | Lignan glycosides |
| 23 | 11 | Neolignans |
| 24 | 2 | Norlignans |
| 25 | 25 | Coumarins<br>Furanocoumarins |
| 26 | 4 | Bibenzyls |
| 27 | 1 | Phenanthrenes |
| 28 | 1 | Dihydrophenanthrenes |
| 29 | 1 | Miscellaneous stilbenoids |
| 30 | 5 | Stilbenoid dimers, stilbenoid oligomers and bis(bibenzyl)s |
| 31 | 1 | Diarylheptanoids |
| 32 | 1 | Zingiber derived compounds |
| 34 | 8 | Linear monoterpenes |
| 35 | 45 | Cyclic monoterpenes |
| 36 | 22 | Iridoids<br>Secoiridoids |
| 37 | 2 | Others |
| 38 | 170 | Farnesenes<br>Germacrenes<br>Bisabolanes<br>Humulenes<br>Eudesmanes<br>Guaianolide<br>Pseudoguaianolide<br>Tutinolide<br>Others<br>Linear diterpenes<br>Apocarotenoids |
| 39 | 3 | Cadinanes |
| 40 | 5 | Abietanes |
| 41 | 43 | Aphidicolane<br>Daphnanes<br>Gibberellins<br>Kaurenes<br>Podocarpanes<br>Tiglianes<br>Others |
| 42 | 3 | Cembrenes |
| 43 | 11 | Clerodanes |
| 44 | 2 | Ginkgolides |
| 45 | 2 | Grayanoids |
| 46 | 5 | Labdanes |
| 47 | 2 | Mesotricyclic diterpenoids |
| 48 | 2 | Pimaranes |
| 49 | 5 | Taxanes |
| 50 | 2 | Liner triterpenes |
| 51 | 38 | Dammarenes<br>Limonoids<br>Lupanes<br>Oleananes<br>Protostanes<br>Ursanes<br>Others |
| 52 | 2 | Hopanes |
| 53 | 1 | Estrane |
| 54 | 2 | Androstane |
| 55 | 1 | Pregnane |
| 56 | 6 | Cardanolide |
| 57 | 5 | Bufanolide |
| 58 | 2 | Others |
| 59 | 8 | Carotenoids |
| 60 | 1 | Polyterpenoids |
| 61 | 5 | Anthrone type |
| 62 | 8 | Anthraquinone type |
| 63 | 3 | alpha-Pyrones |
| 64 | 2 | Monocyclic gamma-pyrones |
| 65 | 2 | Naphthopyrones |

| | | |
|---|---|---|
| 66 | 3 | Cannabinoids |
| 67 | 6 | Phloroglucinols |
| 68 | 6 | Saturated fatty acids<br>Unsaturated fatty acids |
| 69 | 5 | Polyynes |
| 70 | 4 | Others |
| 71 | 5 | Betacyanins<br>Betaxanthins<br>Others |
| 72 | 4 | Cyanogenic glucosides derived from phenylalanine<br>Cyanogenic glucosides derived from thyrosine |
| 73 | 4 | Cyanogenic glucosides derived from valine or isoleucine<br>Cyanogenic glucosides derived from leucine<br>Others |
| 74 | 1 | Cyanogenic glucosides derived from nonprotein amino acid |
| 75 | 1 | Cyanogenic glucosides derived from other origin |
| 76 | 1 | Glucosinolates derived from methionine<br>Glucosinolates derived from leucine<br>Glucosinolates derived from isoleucine<br>Glucosinolates derived from valine<br>Glucosinolates derived from alanine |
| 77 | 1 | Glucosinolates derived from phenylalanine<br>Glucosinolates derived from tyrosine<br>Glucosinolates derived from tryptophan |
| 78 | 4 | Amines |
| 79 | 2 | Amides |
| 80 | 5 | alpha-Naphthoquinones |
| 81 | 7 | Galloyl derivatives<br>Ellagitannins |
| 82 | 1 | Gallotannins |
| 83 | 1 | Miscellaneous galloyl derivatives |
| 84 | 5 | Others |
